# Repetition plasticity in primary auditory cortex occurs across long timescales for spectrotemporally randomized pure-tones

**DOI:** 10.1101/2023.04.26.538446

**Authors:** Nasiru K. Gill, Nikolas A. Francis

## Abstract

Repetition plasticity is a ubiquitous property of sensory systems in which repetitive sensation causes either a decrease (“repetition suppression”, i.e. “adaptation”) or increase (“repetition enhancement”, i.e. “facilitation”) in the amplitude of neural responses. Timescales of repetition plasticity for sensory neurons typically span milliseconds to tens of seconds, with longer durations for cortical vs subcortical regions. Here, we used 2-photon (2P) imaging to study repetition plasticity in mouse primary auditory cortex (A1) layer 2/3 (L2/3) during the presentation of spectrotemporally randomized pure-tone frequencies. Our study revealed subpopulations of neurons with repetition plasticity for equiprobable frequencies spaced minutes apart over a 20-minute period. We found both repetition suppression and enhancement in individual neurons and on average across populations. Each neuron tended to show repetition plasticity for 1-2 pure-tone frequencies near the neuron’s best frequency. Moreover, we found correlated changes in neural response amplitude and latency across stimulus repetitions. Together, our results highlight cortical specialization for pattern recognition over long timescales in complex acoustic sequences.

## Main Text

The dynamics of neural responses to repetitive sensation, i.e., “repetition plasticity”, are influenced by the irregularity and duration of time-intervals between sensory events^1-14^. “Repetition suppression,” i.e., “adaptation,” is a decrease in neuronal responsiveness to repeated sensory input and is thought of as a mechanism for efficient coding of sensory information. “Repetition enhancement,” i.e., “facilitation”, is an increase in neuronal responsiveness to repeated sensory input and is believed to reflect neural predictions about the reoccurrence of sensory events. In the auditory system, repetition plasticity has been observed in the inferior colliculus^7,15^, medial geniculate body^13^, and auditory cortex^1,4,5,8-12,14,16-19^. It has also been observed in visual^3,5^ and somatosensory cortices^5,20^. Mechanisms such as synaptic depression and interneuron inhibition are believed to play a role in both cortical and subcortical repetition plasticity^2,3,6,13,17-19,21-24^.

The spectrotemporal context of a sound is an important factor in repetition plasticity. For example, in mice the magnitude of stimulus-specific adaptation in auditory cortex depends on the proportion of “standard” vs. “deviant” pure-tone frequencies presented during an experiment^4,5,8-12,14,17,19^. Repetition plasticity in auditory cortex has been observed on timescales ranging from tens of milliseconds to tens of seconds^5,8-12,19,25^, in comparison to subcortical structures where repetition plasticity occurs on a shorter timescale, typically on the order of tens to hundreds of milliseconds^1,13,25^. Thus, auditory cortex may be specialized to encode global information about irregular and slow acoustic sequences^1,6,23,25^.

To investigate cortical specialization for long timescales in repetition plasticity, we used 2P imaging to study how neurons in mouse A1 L2/3 respond to pure-tone frequencies whose repetition occurred over minutes-long intervals and with low predictability due to equiprobable stimulus statistics (figure 1). We recorded auditory responses to spectrotemporally randomized pure-tones in 874 A1 L2/3 neurons across 26 experiments in 6 awake Thy1-GCaMP6s mice^26^ (figure 1e-k). We found repetition plasticity that progressed slowly over an approximately 20-minute experiment in 48% of the total neuronal population (figure 2). Repetition plasticity occurred on average across subpopulations within a single experiment (middle rows in a1 and a2), and for individual neurons (bottom rows in a1 and a2). Note that while each trace in figure 2a is evenly spaced on the panels, during experiments the frequency-repetition interval was randomized, averaging at 1 ± 0.9 minute standard deviations (SDs) between each of the 20 repetitions spanning the approximately 20-minute experiment (figure 1a-d; top of figure 2).

**Figure 1.**
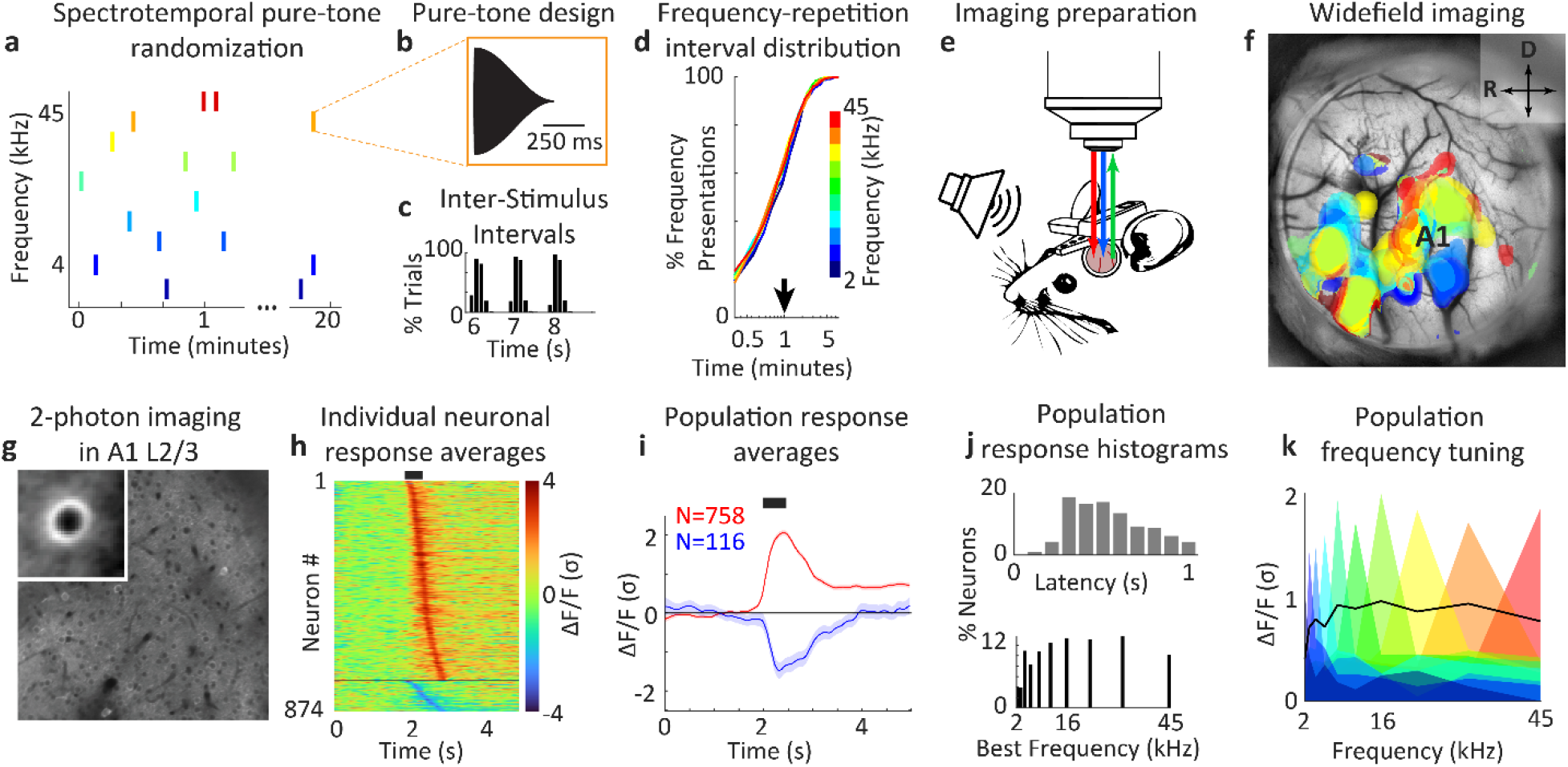
2-photon (2P) imaging of neural activity in primary auditory cortex (A1) layer 2/3 (L2/3) in response to pure-tones. **a**. Stimulus randomization over a 20-minute period. **b**. Stimulus design. **c**. Inter-stimulus Interval histogram. **d**. Frequency-repetition interval cumulative distribution. The distribution for each possible frequency is shown and color-coded. Note that the distributions are overlapping, indicating similar randomizations (1 ± 0.9 minute standard deviations (SDs)). **e**. Experimental setup. A 940 nm 2P laser (red) and 470 nm LED (blue) were used to image neuronal activity (green) in A1 L2/3 of Thy1-GCaMP6s transgenic mice during pure-tone presentations. **f**. Widefield imaging of auditory cortex. Frequency-dependent response amplitude was used to color-code pixels in cortical space, i.e., to find ‘tonotopy’. A1 was identified by the rostrocaudal gradient of tonotopy. **g**. Example 2P imaging field of view (FOV). The inset figure shows the average cell in the FOV. **h**. Heatmap of individual neuronal responses to pure-tones, averaged across all stimuli for each neuron. Fluorescence values (ΔF/F) were normalized by the standard deviation of response amplitude (σ) for each neuron. **i**. Population-averaged response traces. Red: positive responses, Blue: negative responses. Shading shows 2 standard errors of the mean (SEMs). **j**. Top panel: Population peak-latency response distribution. The average peak-latency response was 0.46 ± 0.23 s (SDs). Bottom panel: Best frequency (BF) histogram. **k**. Population-average frequency tuning curves (FTCs). The black line shows the mean tuning curve across the recorded population.

**Figure 2.**
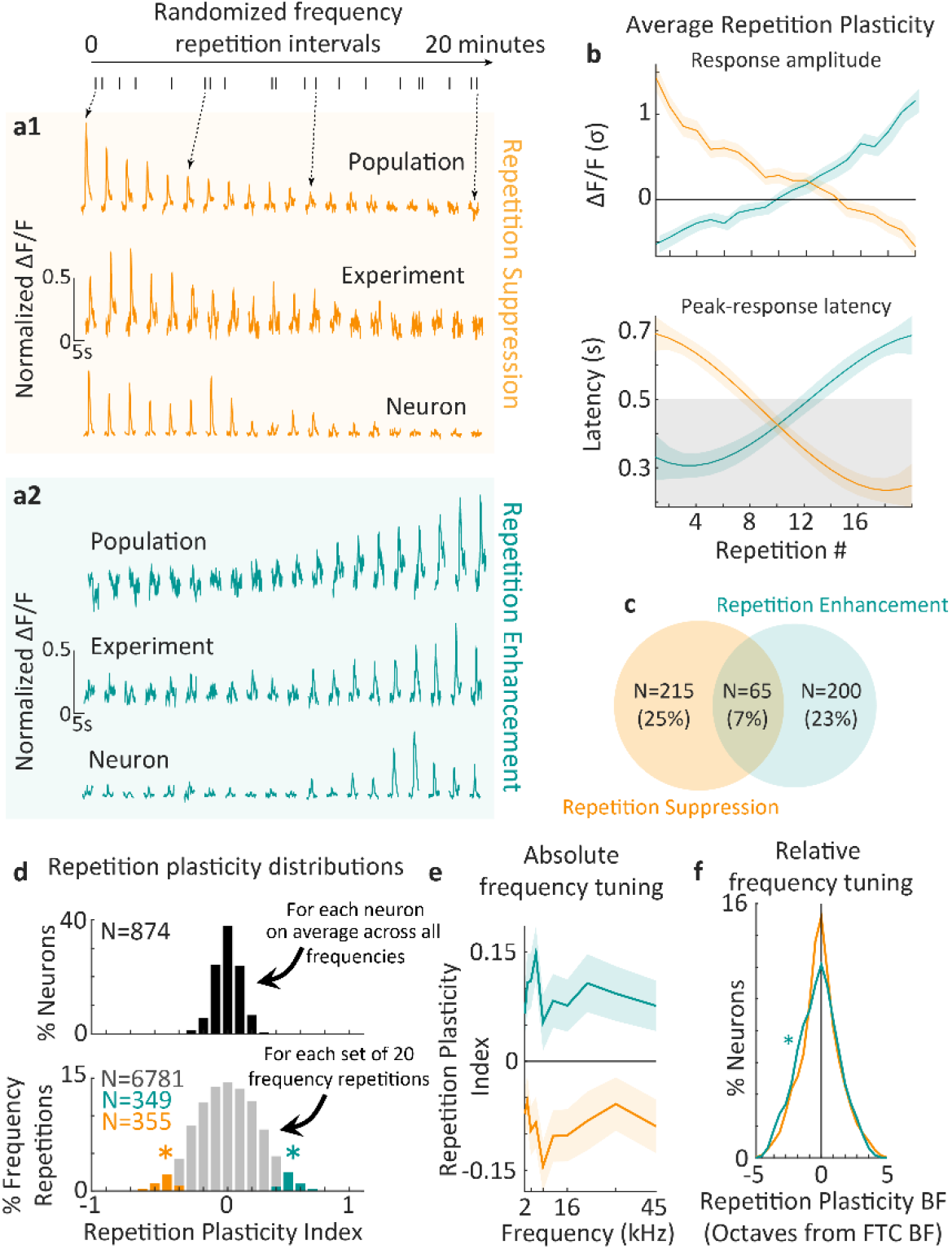
Repetition plasticity (RP) in A1 L2/3. **a1**,**a2**. RP was observed, (1) on average for the population (top rows in a1 and a2), (2) on average within a single experiment (middle rows in a1 and a2), and (3) for individual neurons (bottom rows in a1 and a2). Each row shows unity normalized ΔF/F activity in response to a repetition of a given pure-tone frequency. Note that the traces shown here are equally spaced in time for ease of visualization, however, as shown at the top of the figure, during experiments frequency-repetition intervals were randomized (average: 1 ± 0.9 minutes SDs) over an approximately 20-minute period. **b**. Population-averaged RP (N=874). The top panel shows the average ΔF/F (σ) amplitude in the 1-second interval following each stimulus presentation during each of 20 pure-tone frequency repetitions. The bottom panel shows the average change in peak-response latency across frequency-repetitions. **c**. Venn diagram of the subpopulation sizes for repetition suppression, repetition enhancement, or both. **d**. RP distributions. The RP index was defined as the correlation coefficient across each set of 20 pure-tone frequency-repetitions. The top panel shows that none of the neurons in our population had significant repetition plasticity on average across all frequency-repetitions (p>0.05). However, the lower panel expands the data in the top panel by showing the histogram of repetition plasticity for each set of 20 frequency-repetitions in each neuron (N=8004). Asterisks indicate significant repetition suppression (N=349) or enhancement (N=355) (p<0.05). Gray bars show repetitions with insignificant RP (N=6781, p>0.05). **e**. Population-averaged RP absolute frequency tuning curves. Significant (p<0.016) RP was found for all pure-tone frequencies and with near-uniform magnitude across frequency. **f**. Distribution of RP BF relative to FTC BF. The RP BF tended to occur near the FTC BF, though the average frequency with the greatest RP tended to be just below BF (−0.25 ± 1.7 octave SDs, p=0.017 and -0.1 ± 1.6 octave SDs, p=0.32, respectively). In all panels, shading shows 2 SEMs.

Figure 1a illustrates the time-course of stimulus presentation in our experiments. Pure-tones were presented at 70 dB SPL and 500 ms in duration, with 5 ms and 495 ms onset and offset ramps, respectively (figure 1b). Inter-stimulus intervals were randomly selected from a trimodal distribution peaking at 6, 7, or 8 s (figure 1c). For each presentation, the pure-tone frequency was randomly selected between 2-45 kHz (10 possible frequencies spaced 0.5 octaves apart) (figure 1d). Each frequency was repeated 20 times during an experiment. Thus, the presentation of each pure-tone frequency was equiprobable and irregularly distributed in time, forming a slow and complex acoustic sequence.

We began our experiments using widefield imaging to localize A1 in each mouse (figure 1e). Figure 1f shows a color-coded mapping of pure-tone frequency selectivity across space in auditory cortex, i.e., “tonotopy,”. A1 is identified by a rostro-caudal gradient of high-to-low frequencies in the posterior region of auditory cortex. Once A1 was localized in each mouse, we then used 2P imaging to record pure-tone responsiveness in populations of individual neurons that were approximately 150 µm below the cortical surface in A1 L2/3 (figure 1g). Figure 1h shows a heatmap of the average stimulus-aligned responses from individual neurons, sorted by peak-response latency. Most neurons (N=758, 87%) had positive responses (increases from the silent baseline before a pure-tone in each presentation). A smaller population (N=116, 13%) had negative responses (decreases from baseline). The average traces across positive and negative response populations are shown in figure 1i. The average peak-latency of neural responses to pure-tones was 460 ms ± 230 ms (SDs) (figure 1j, top panel), partly predicated by stimulus-locked responsiveness (figure 1h-j) and the sluggish rise-time of the GCaMP6s fluorescence indicator^26^.

To maximize the number of individual neurons across experiments, and to ensure that the neural population under study did not have a frequency-selectivity bias, we chose a different 2P field of view within A1 L2/3 for every experiment. For each neuron, we calculated the average magnitude of its response to each of the 10 pure-tone frequencies to create a frequency tuning curve (FTC). We then found the frequency with the largest response, i.e., the “best frequency,” (BF) from the FTC (figure 1j, bottom panel). FTCs from neurons with the same BF were averaged together and are plotted in figure 1k, color-coded by BF. Our results show that we imaged neurons with BFs evenly distributed across the range of pure-tone frequencies, indicating that the mice had healthy hearing in the tested frequency range. Given our widefield tonotopy results, combined with stimulus-locked response latencies and well-defined FTCs, it is likely that we successfully targeted auditory neurons in A1 L2/3 during 2P imaging.

It is important to note that one might not expect repetition plasticity to occur in our experiments because of extensive spectrotemporal pure-tone randomization, and indeed, we did not find repetition plasticity on average across all frequency-repetitions for a given neuron (p>0.05) (figure 2d, top panel). However, upon finer parcellation of the data into individual frequency-repetitions (N=8004 repetitions), we found subpopulations of neurons with repetition suppression (N=215), repetition enhancement (N=200), or both (N=65) (Figure 2c).

Repetition plasticity occurred for only a subset of frequencies in each neuron. The average number of frequencies with repetition plasticity per neuron was 1.3 +/-0.7 SDs. Figure 2e shows that the frequency tuning curves for both repetition suppression and enhancement tended to be uniform, and thus occurred across the range of presented frequencies. In contrast, figure 2f shows that the frequency with the greatest (“best”) repetition plasticity tended to occur near the neuron’s BF, though the distributions of best repetition enhancement and suppression frequencies were skewed below the BF (averages: -0.25 ± 1.7 octave SDs, p=0.017 and -0.1 ± 1.6 octave SDs, p=0.32, respectively).

Since cortical response amplitudes and latencies are both state- and stimulus-dependent in auditory cortex^7,27,28^, here we quantified the effect of slow and irregular stimulus repetition on neuronal peak-response latencies. Consistent with the opposing amplitude changes we observed for repetition suppression vs enhancement, we found that repetition suppression neurons tended to begin with short response latencies that became longer over repetitions, and vice versa for repetition enhancement neurons (Figure 2b, bottom panel). Thus, we find that repetition plasticity slowly and monotonically changes both the timing and amplitude of neural responses to sound.

Here we describe subpopulations of neurons in A1 L2/3 that encode stimulus repetition over a period lasting tens of minutes, despite extensive randomization in stimulus design. This long timescale of repetition plasticity reflects the importance of cortical processing for pattern recognition in slow and complex acoustic sequences. It may be important that our mice were naïve to hearing pure-tones at the start of the experiment. Thus, it is possible that the relative stimulus novelty drew their attention to the pure-tones, which may have affected neural activity in A1^27,28^. It remains to be seen how mechanisms such as synaptic depression and interneuron inhibition—processes typically associated with timescales limited to hundreds of milliseconds—might sustain repetition plasticity over minutes, but perhaps their involvement in long-range recurrent network activity plays a role^6,22-25^.

## Acknowledgements

Supported by NIH R21DC017829 (NAF).

## Methods

### Experimental model and subject details

All procedures were approved by the University of Maryland Institutional Animal Care and Use Committee. We used N=6 mice (3 female, 3 male) F1 offspring of CBA/CaJ mice (The Jackson Laboratory; stock #000654) crossed with transgenic C57BL/6J-Tg(Thy1GCaMP6s)GP4.3Dkim/J mice^26^ (The Jackson Laboratory; stock #024275), 1.5-7 months old, in 26 total experiments. We used the F1 generation of the crossed mice because they have healthy hearing at least 1 year into adulthood^29^. Mice were housed under a reversed 12 h-light/12 h-dark light cycle.

### Stimulus Design and Presentation

We presented awake mice with 70 dB SPL pure-tones from a free-field speaker (Figure 1). Each pure-tone was 500 ms in duration, with 5 ms and 495 ms raised-cosine attack and decay ramps, respectively. The frequency of each pure-tone was randomly selected from 10 equiprobable values (2-45 kHz, 2 tones per octave). Each frequency was repeated 20 times per experiment, with a frequency-repetition interval of 1 +/-0.9 minute standard deviations (SDs). Inter-stimulus intervals were randomized according to a tri-modal distribution (peaks at 6, 7, and 8 s) across the duration of each experiment, lasting approximately 20 minutes.

### Chronic window implantation

Mice were given an intraperitoneal injection of dexamethasone (5mg/kg) at least 1 hour prior to surgery to prevent inflammation and edema. Mice were deeply anesthetized using isoflurane (5% induction, 0.5-2% for maintenance) and given a subcutaneous injection of cefazolin (500mg/kg). Internal body temperature was maintained at 37.5 C using a feedback-controlled heating blanket. Scalp fur was trimmed using scissors and any remaining fur was removed using Nair. The scalp was disinfected with alternating swabs of 70% ethanol and betadine. A patch of skin over the temporal bone was removed and the underlying bone cleared of connective tissue using a scalpel. The temporal muscle was detached from the skull, and the skull was cleaned and dried. A thin layer of cyanoacrylate glue (VetBond) was applied to the exposed skull surface and a 3D printed stainless steel head-plate was affixed to the midline of the skull. Dental cement (C&B Metabond) was used to cover the entire head-plate. A circular craniotomy (3 mm diameter) was made over auditory cortex where the chronic imaging window was then implanted. The window was either of a stack of two 3 mm diameter coverslips or a 3.2 mm diameter, 1 mm thick uncoated sapphire window (Edmund Optics), glued with optical adhesive (Norland 61) to a 5 mm diameter coverslip. The space between the glass and the skull was sealed with a silicone elastomer (Kwik-Sil). The edges of the glass and the skull were then sealed with dental cement. Finally, the entire implant except for the imaging window was coated with black dental cement created by mixing methyl methacrylate with iron oxide powder to reduce optical reflections. Meloxicam (0.5mg/kg) was given subcutaneously as a post-operative analgesic. Animals were allowed to recover for 2 weeks prior to imaging experiments.

### Widefield imaging

Awake mice were placed into a 3D-printed plastic tube and head-restraint system. Blue excitation light was shone by an LED (470 nm) through an excitation filter (470 nm) and directed into the cranial window. Emitted fluorescence (F) from neurons in Thy1-GCaMP6s mice was collected through a 4x objective (Thorlabs), passed through a long-pass filter (cutoff: 505 nm), followed by a bandpass emission filter (531 nm) attached to a pco.panda 4.2 CMOS camera. Images were acquired using Micro-manager software. After acquiring an image of the cortical surface, the focal plane was advanced to approximately 500 μm below the surface.

Our goal was to visualize primary auditory cortex (A1) by identifying a rostro-caudal tonotopic gradient in the posterior region of auditory cortex. To visualize tonotopy, pure-tones were presented from a free field speaker, as described above. Widefield images were acquired at a 30 Hz rate and 256×288 pixels. Using Matlab software (The Mathworks), image sequences for each tone frequency were averaged and processed with a homomorphic (contrast) filter to extract reflectance^27^. For each pixel, ΔF/F traces were calculated by finding the average F taken from the silent baseline period before a pure-tone presentation, subtracting that value from subsequent time-points until 3s after the pure-tone, then dividing all time-points by the baseline F. To visualize auditory responses, we kept traces with ΔF/F within 90% of the maximum response in the pixel-wise grand-average of ΔF/F (i.e., ΔF/F_90_). Pixel-wise tonotopic frequencies were taken as the median frequency of the set of tones corresponding to the ΔF/F_90_ traces (figure 1f).

### 2-photon imaging

After visualizing A1 tonotopic maps using widefield imaging, recording sites were selected for 2-photon (2P) imaging in A1 layer 2/3 (L2/3) for each mouse. Our 2P recording sites were chosen at various regions across A1 (figure 1f). We used a scanning microscope (Bergamo II series, Thorlabs) coupled to a pulsed femtosecond 2-photon laser with dispersion precompensation (Coherent Chameleon Discovery NX TPC). The microscope was controlled by ThorImage software. The laser was tuned to λ = 940 nm to excite GCaMP6s. Fluorescence signals were collected through a 16× 0.8 NA microscope objective (Nikon). Emitted photons were directed through a 525 nm (green) bandpass filter onto a GaAsP photomultiplier tube. The field of view was 411 × 411 µm. Imaging frames of 512×512 pixels (0.8 µm per pixel) were acquired at 30 Hz by bidirectional scanning of an 8 kHz resonant scanner. Laser power was set to approximately 70 mW, measured at the objective. During experiments, the objective’s focal plane was lowered into L 2/3 (∼150 µm below the surface) before imaging neuronal responses to pure-tones.

After 2P experiments, all images were processed using Matlab^27^. Image motion was corrected using the TurboReg plug-in for MIJI (i.e., FIJI for Matlab). Figure 1g shows the average of registered images for GCaMP6s images. After manually selecting the centers of cell bodies, a ring-like region of interest (ROI) was cropped around the cell center. Overlapping ROI pixels (due to neighboring neurons) were excluded from analysis. For each labeled neuron, a raw fluorescence signal over time was extracted from somatic ROIs. Pixels within the ROI were averaged to create individual neuron fluorescence traces, F_C_(t), for each trial of the experiment. Neuropil fluorescence was estimated for each cellular ROI using an additional ring-shaped ROI, which began 3 pixels from the somatic ROI. Pixels from the new ROI were averaged to obtain neuropil fluorescence traces, F_N_(t), for the same time-period as the individual neuron fluorescence traces. Pixels from regions with overlapping neuropil and cellular ROIs were removed from neuropil ROIs. Neuropil-corrected cellular fluorescence was calculated as 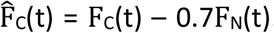. Only cells with positive values obtained from averaging 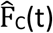 across time were kept for analysis, since negative values may indicate neuropil contamination. ΔF/F was calculated from 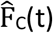, for each neuron, by finding the average F taken from the silent baseline period before a pure-tone presentation, subtracting that value from subsequent time-points until 3s after the pure-tone, then dividing all time-points by the baseline F.

### Quantifying repetition plasticity

We quantified the modulation of cortical activity by repetitive sensation, i.e., “response plasticity”, by (1) taking the average ΔF/F in the 1-second interval following each stimulus presentation, (2) concatenating averaged ΔF/F values to form a sequence of 20 values taken from each of the 20 frequency-repetitions, and (3) calculating the correlation coefficient across a given set of 20 values. We refer to the resulting correlation coefficient as the, “Repetition plasticity index”. For the same sequence of 20 stimulus presentations, we quantified response latencies by finding the time of the peak ΔF/F response for each stimulus, then fitting a cubic polynomial to each set of 20 response latencies.

### Statistical analysis

Statistical comparisons were performed using a non-parametric bootstrap test with 10000 iterations. All mean values are reported with either standard deviations (SDs) or standard errors of the mean (SEMs).

